# Gastrocnemius muscle remodeling explains functional deficits three months following Achilles tendon rupture

**DOI:** 10.1101/585505

**Authors:** Todd J. Hullfish, Kathryn M. O’Connor, Josh R. Baxter

**Author notes:** Corresponding author: Josh R. Baxter, PhD. **Mailing address:** 3737 Market Street, Suite 702, Philadelphia, PA, USA 19104.

## Abstract

Plantarflexor functional deficits are associated with poor outcomes in patients following Achilles tendon rupture. In this longitudinal study, we analyzed the fascicle length and pennation angle of the medial gastrocnemius muscle and the length of the Achilles tendon using ultrasound imaging. To determine the relationship between muscle remodeling and functional deficits measured at 3 months after injury, we correlated the reduction in fascicle length and increase in pennation angle with peak torque measured during isometric plantarflexor contractions and peak power measured during isokinetic plantarflexor contractions. We found that the medial gastrocnemius underwent an immediate change in structure, characterized by decreased length and increased pennation of the muscle fascicles. This decrease in fascicle length was coupled with an increase in tendon length. These changes in muscle-tendon structure persisted throughout the first three months following rupture. Deficits in peak plantarflexor power were moderately correlated with decreased fascicle length at 120 degrees per second (R^2^ = 0.424, *P* = 0.057) and strongly correlated with decreased fascicle length at 210 degrees per second (R^2^ = 0.737, *P* = 0.003). However, increases in pennation angle did not explain functional deficits. These findings suggest that muscle-tendon structure is detrimentally affected following Achilles tendon rupture. Plantarflexor power deficits are positively correlated with the magnitude of reductions in fascicle length. Preserving muscle structure following Achilles tendon rupture should be a clinical priority to maintain patient function.

## INTRODUCTION

Achilles tendon ruptures have increased by a factor of 10 in the past three decades, mostly affecting physically-active males.^1^ Traditionally, these injuries have been treated as isolated tendon injuries by surgically repairing ruptured tissue to re-oppose the ‘mop-like’ ends of the tendon in order to provide temporary stability throughout healing. However, in the past decade, surgeons in Scandinavia have opted to treat three out of four patients non-surgically.^2^ Although rerupture rates are below 5%, regardless of surgical or non-surgical treatment,^3^ functional deficits persist in two out of three patients.^4^

Plantarflexor work and power partially govern ambulatory function. Previous studies have documented tendon elongation following Achilles tendon ruptures,^5–7^ which has been strongly correlated with decreased single-leg heel raise height.^6^ However, tendon is not a contractile tissue and cannot perform the positive work necessary to produce human motion. Instead, plantarflexor work and power is associated with muscle shortening dynamics, where adults with longer gastrocnemius fascicles generate greater plantarflexor work and power.^8^ Single-leg heel raise height is positively correlated with resting gastrocnemius length^9^ and active shortenng.^10^ Following Achilles tendon ruptures, patients have compromised plantarflexor function and compensate by increasing the work done by the proximal knee joint.^11^

Gastrocnemius muscle remodeling occurs following Achilles tendon ruptures,^12,13^ which appears to be in response to a sudden loss of muscle-tendon tension. Acute Achilles tendon ruptures have been shown to elicit shorter and more pennate gastrocnemius muscle fascicles at rest.^12^ This muscle reconfiguration persists through at least 4 weeks when the Achilles tendon rupture is treated non-surgically.^12^ Additionally, we reported that long-term functional deficits were explained by permanent reductions in fascicle length and increases in pennation angle in a case study.^14^ However, the effects of Achilles tendon ruptures on early muscle structure and function have not yet been established. Understanding this muscle remodeling timeline in the first 3 months following injury and how muscle structure impacts patient function would provide critical new insight into the efficacy of non-surgical treatment.

The purpose of this study was to link plantarflexor work and power with gastrocnemius muscle remodeling and tendon elongation 3 months following acute Achilles tendon ruptures. To do this, we measured resting muscle structure at each clinical visit from initial presentation to 3 months and plantarflexor kinetics using an isokinetic dynamometer 3 months after injury. We hypothesized that the magnitude of muscle remodeling would correlate with the magnitude of functional deficits. Based on our experimental measurements in healthy adults^8^ and computational models,^9^ we expected that fascicle length changes would be a stronger correlate of plantarflexor functional deficits than pennation angle. Additionally, we hypothesized that our previous measurements of muscle remodeling – characterized by shorter and more pennate gastrocnemius muscle fascicles – would persist through 3 months.

## METHODS

### Study Design

Ten adults (9 males, age: 44 ± 11 years, BMI: 28.7 ± 6.3) who suffered acute Achilles Tendon ruptures and were treated nonsurgically by a fellowship trained foot and ankle surgeon and underwent accelerated rehabilitation^15^ were enrolled in this IRB approved study. Patients provided written informed consent prior to participation. All patients were recruited from the clinics of the Department of Orthopaedic Surgery at the Penn Medicine and met several inclusion criteria: patient was between 18 and 65 years old and elected to be treated nonsurgically for acute Achilles tendon rupture within 2 weeks of injury. Patients were not enrolled in this study if they were had excessive weight (BMI > 50) or concomitant lower extremity injuries. We measured the medial gastrocnemius of the unaffected limb at the initial visit and longitudinally tracked medial gastrocnemius structure of the affected limb at each of six clinical visits (**Figure 1**): time of injury when patients were placed in a cast in maximal plantarflexion (week 0); time of cast removal when patients were transitioned into partial weight bearing in a walking boot with crutches (week 2); transition from partial to full weight bearing with crutches (week 4); transition to full weight bearing without crutches (week 4-6); transition out of walking boot (week 10); and when patients were cleared to begin using the injured limb functionally (week 14). In addition to measurements of muscle structure, plantarflexor function of both limbs was measured during two of the clinical visits to assess the magnitude of deficits 3 months after injury.

**Figure 1.**
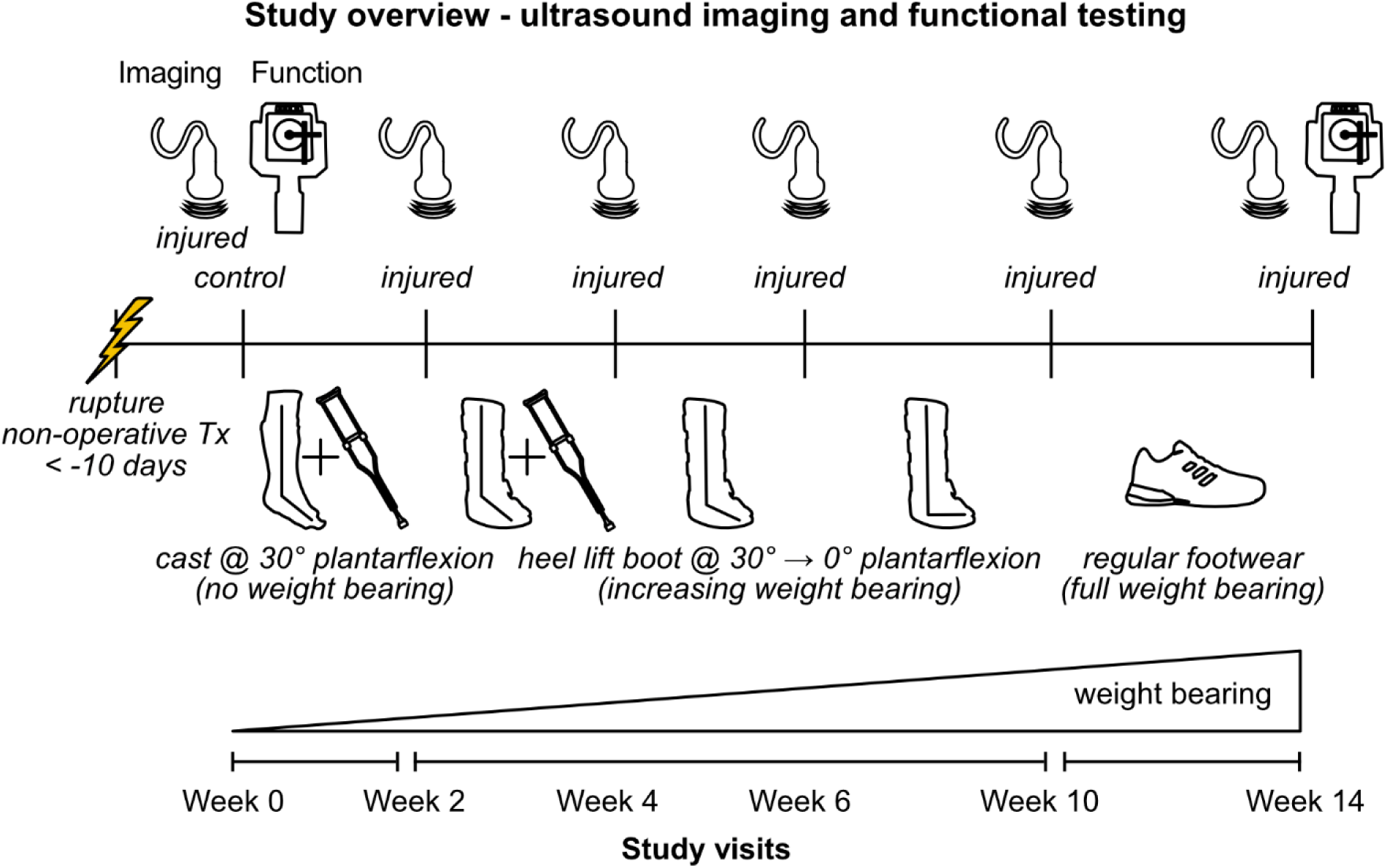
Muscle-tendon structure and function was assessed during the first three months of treatment at each clinical visit. Ultrasound images were acquired to quantify medial gastrocnemius fascicle length and pennation and Achilles tendon length. Isokinetic dynamometry was performed at three months on the injured limb and within the first two weeks following initial presentation of injury on the contralateral limb (*control*). All patients were treated nonsurgically and were prescribed the same progressive weight bearing protocol where patients were first placed in a plantarflexed cast and used crutches for 2 weeks before transitioning to a walking boot. Once in the walking boot, patients decreased the amount of plantarflexion the foot was held in before transitioning to normal footwear around week 10.

### Muscle-Tendon Structure

To determine how muscle and tendon structure changes following injury, we acquired longitudinal B-mode ultrasound images of the medial gastrocnemius and Achilles tendon of both the affected and contralateral limb at each clinical visit. Using an 8 MHz transducer with a 6 cm scanning width (LV7.5/60/128Z-2, SmartUs, TELEMED) and constant scanning parameters for all patients and imaging sessions (Dynamic Range: 72 dB; Frequency: 8 MHz; Gain: 47 dB), we acquired continuous images of the mid muscle belly while patients lay prone on a treatment table. To ensure reliable imaging, a single investigator positioned the probe using established imaging guidelines.^16^ Briefly, we positioned the probe on the central portion of the muscle belly halfway between the muscle-tendon junction and the crease of the knee where the deep and superficial aponeuroses of the muscle were seen to be parallel. We then rotated the probe to ensure that the muscle fascicles were aligned with the imaging plane. To ensure consistent tendon loading during each visit, we first identified patient resting angle^17^ and then shifted the patient proximally on the treatment table to support the weight of the foot and not load the healing tendon. This was of particular focus during the first 8 weeks of healing because patients were treated nonsurgically and accidental loading could cause tendon elongation. To measure Achilles tendon length, we measured the distance from the muscle-tendon junction of the medial gastrocnemius to the calcaneal insertion using a 15 MHz transducer with a 4 cm scanning width (L15-7L40H-5, SmartUs, TELEMED), which we measured with a reflective marker and 12-camera motion capture system (Raptor Series, Motion Analysis Corporation, Rohnert Park, CA, USA). Using a custom printed probe mold with reflective markers, we experimentally determined the rigid body transform to convert ultrasound image data to the lab coordinate system. Our internal validation efforts confirmed that this ultrasound technique was accurate to under 2 mm.

We quantified muscle-tendon structure of the medial gastrocnemius muscle using established techniques. Using custom written software (MATLAB 2018a, The MathWorks, Inc., Natick, MA), we anonymized and randomized each image to ensure that the investigator performing the analysis could not be biased. We measured medial gastrocnemius structure as the length and pennation of a centrally positioned fascicle in the ultrasound image. For each muscle image, the investigator identified the deep and superficial aponeuroses as well as a single fascicle (**Figure 2**). We defined fascicle length as the point to point distance between the insertions into the aponeuroses and pennation angle as the angle between the fascicle and the deep aponeurosis. The same investigator identified the muscle-tendon junction of the medial gastrocnemius muscle and calculated the distance to the calcaneal insertion. To confirm the repeatability of this measurement, we calculated tendon length of the control limb at each visit and found our measurements to be accurate to within 0.8% on average and 1.9% at most compared to the average tendon length.

**Figure 2.**
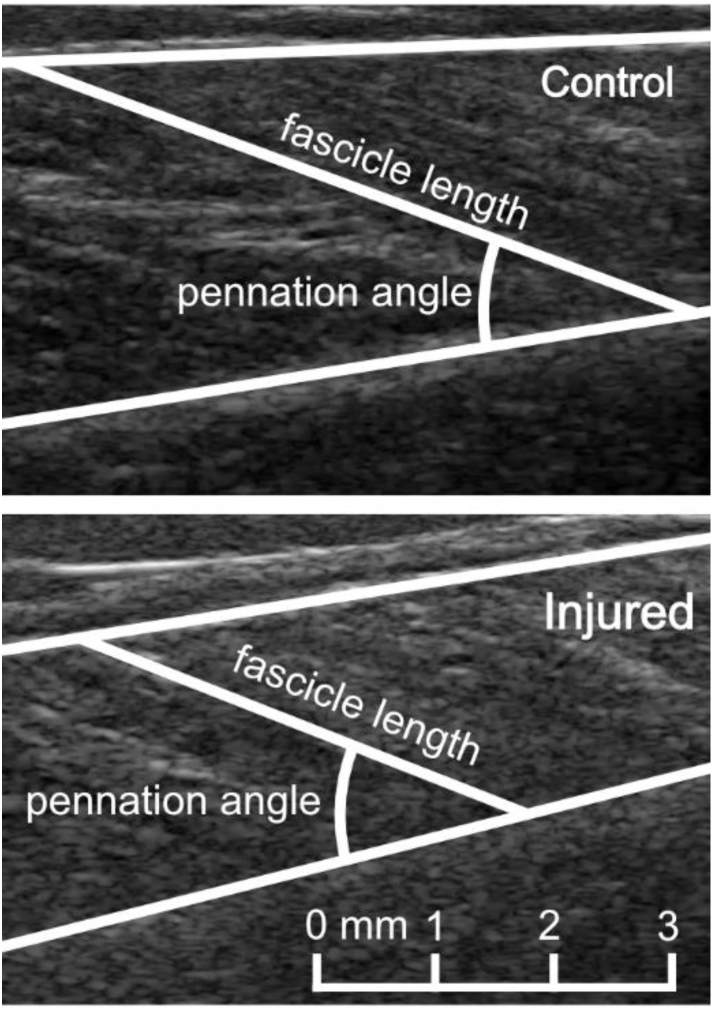
Ultrasound images of the medial gastrocnemius muscles on the control (*top*) and injured (*bottom*) limbs were acquired and analyzed to calculate fascicle length and pennation angle.

### Functional Assessment

To determine the effects of muscle remodeling on plantarflexor function 3 months following initial injury, we implemented a combined isokinetic dynamometry and ultrasound imaging framework. We positioned patients prone on a treatment table that was rigidly attached to a dynamometer (System 4, Biodex) with their knees fully extended. The spindle of the dynamometer was aligned with the medial malleolus of the involved ankle and the foot was secured to a foot plate with two straps. To prevent movement on the treatment table during maximal contractions, patients held onto handles located on the sides of the treatment table. The same ultrasound probe used to image resting muscle structure was secured over the mid muscle belly of the medial gastrocnemius with a custom made cast. Continuous B-mode ultrasound images of muscle were acquired at 30 Hz. Dynamometer position, velocity, and torque and an ultrasound syncing pulse were synchronously acquired at 1kHz.

Prior to testing, we measured ankle range of motion by instructing each patient to rotate their foot into as much dorsi-and plantarflexion as they were able voluntarily. The investigator then manually positioned the ankle in neutral and the absolute position of the spindle in this position was used as an offset to determine the position of the ankle for all trials. To account for any differences in ankle position between the passive imaging and functional protocols, measurements of muscle structure were repeated while the dynamometer passively rotated the ankle through the patient specific range of motion. The footplate rotated at 10 degrees per second and we measured fascicle length and pennation throughout the range of motion using an automatic tracking algorithm.^18^

After establishing ankle range of motion and muscle structure, patients performed a series of maximal isometric and isokinetic plantarflexion contractions. Patients performed these isometric contractions with a neutrally-aligned ankle and isokinetic contractions at 30, 120, and 210 degrees per second throughout the entire range of ankle motion. Patients were instructed to push against the foot plate as hard and as fast as they could. Verbal encouragement was provided while pushing and visual feedback of the torque generated was displayed after completing each contraction. Patients performed 3-4 contractions per condition and were given time to rest between conditions to avoid fatigue. This protocol was performed for the unaffected limb as close to time of injury as possible to establish baseline function (week 0 or week 2). The protocol was repeated for the affected limb at the 3-month clinical visit when patients were cleared by their treating physician.

### Statistical Analysis

To test our hypothesis that muscle remodeling persists through three months, we compared fascicle length and pennation angle of the injured limb at each clinical visit to the control limb measured during the initial visit. Because this hypothesis was directional, we used one-tailed paired t-tests and calculated the Cohen’s effect size (*d*) and percent differences from the control limb. We compared tendon length and muscle structure between control data measured at week 0 and the injured side at each visit from week 0 to 10 with both feet supported in the resting angle. When loading restrictions were removed at week 14, we compared fascicle length and pennation at 15 degrees plantarflexion, 0 degrees neutral, and 15 degrees dorsiflexion.

To test our hypothesis that the magnitude of muscle remodeling following an acute Achilles tendon rupture would correlate with the magnitude of functional deficits observed, we performed univariate linear regressions to determine the relationship between differences in peak torque and changes in fascicle length and pennation. The effect sizes of these regression analyses were calculated using the coefficient of determination (R^2^) with values ranging from 0 to 1. These values indicate the strength of correlation in terms of negligible (0.00-0.04), weak (0.04-0.25), moderate (0.25-0.64), and strong (0.64-1.00).^19^ Peak torques for each condition were compared between the baseline and 3-month time point and absolute differences were calculated to be used in statistical analysis. We also performed paired one-way t-tests to test if plantarflexor torque and power was reduced 3 months following initial injury.

## RESULTS

Passive Gastrocnemius muscle structure and Achilles tendon length following an acute Achilles tendon rupture differed with the healthy-contralateral limb throughout the first ten weeks following injury (**Figure 3**). Fascicle length was 14% shorter (d = 1.2, P ≤ 0.019), pennation angle was 24% greater (d = 1.2, P ≤ 0.018), and tendon length was 8% longer (d = 1.6, P ≤ 0.005) at the presentation of injury (week 0). Differences in fascicle length persisted throughout the first 10 weeks (d ≥ 0.8) but returned to within 6% of the healthy-contralateral limb. Pennation angle differed for the first 6 weeks and returned to within 3% of the contralateral limb by week 10. Differences in tendon length persisted throughout the first 10 weeks (d ≥ 1.0, P ≤ 0.005). Additionally, passively loaded muscle structure of the injured limb at 3 months differed from the healthy contralateral limb across all conditions tested (**Figure 4**). Fascicle length was 13-19% shorter (P ≤ 0.003) in plantarflexed, neutral, and dorsiflexed ankle positions. Similarly, pennation angle was increased by 19% in dorsiflexion and neutral alignment (*P* = 0.047), but these differences became not statistically different in plantarflexion (16% increase, *P* = 0.62).

**Figure 3.**
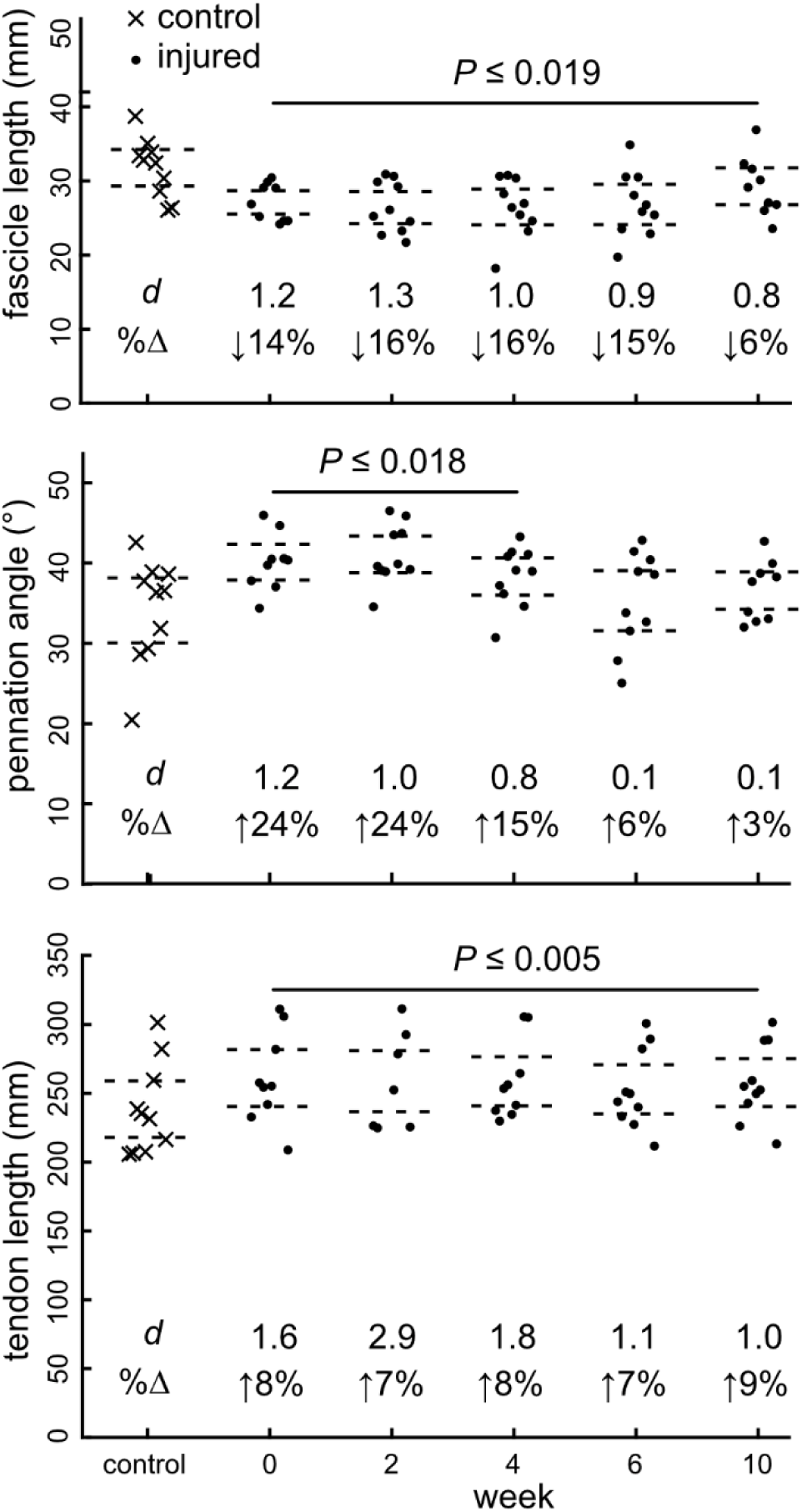
Muscle and tendon structure differed following Achilles tendon rupture compared to the contralateral side when imaged with the foot fully supported in plantarflexion. Fascicle length (*top*) was shorter following rupture and remained shorter thru the first 10 weeks. Conversely, pennation angle (*middle*) increased following the injury but returned to the contralateral limb values at week 6. Tendon length (*bottom*) increased following the injury and remained consistently elongated through 10 weeks. Effect sizes (Cohen’s d) and percent differences (%Δ) are reported in each subfigure.

**Figure 4.**
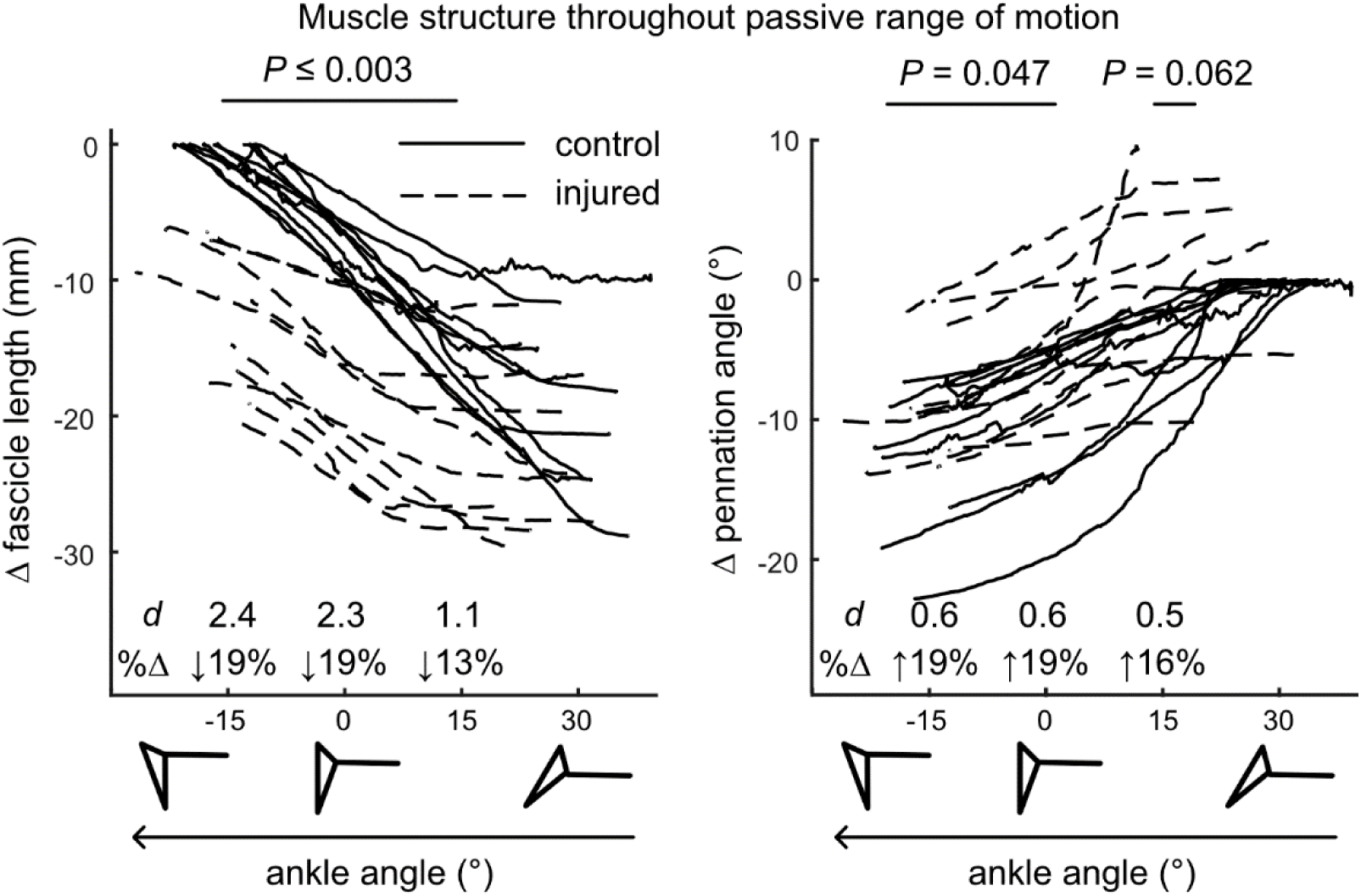
Medial gastrocnemius structure of the injured side (*dashed line*) differed throughout the ankle range of motion compared to the control side (*solid line*) when the foot was passively moved from peak plantarflexion to peak dorsiflexion at 10 degrees per second measured 3 months following rupture. Fascicle length (*left*) was shorter throughout the range of motion (−15, 0, and 15 degrees) and had large effect sizes (Cohen’s *d* > 1.1) compared to the control muscle. Similarly, pennation angle (*right*) was greater throughout the range of motion but the effect sizes were less strong (Cohen’s *d* ∼ 0.6) and the statistical tests yielded *P*-values around our *a priori* threshold of 0.05. The Y-axes are corrected for by the peak values of fascicle length and pennation angle of the control sides to demonstrate the differences in structure caused by tendon rupture.

Changes in isokinetic power (**Figure 5**) were moderately and strongly correlated with changes in fascicle length at 120 degrees per second (R^2^ = 0.424, P = 0.057) and 210 degrees per second (R^2^ = 0.737, P = 0.003). There was no correlation between change in fascicle length and isometric torque or isokinetic power at 30 degrees per second. Changes in pennation angle and tendon length were not correlated with change in peak isometric torque or isokinetic power. Plantarflexion function of the injured limb at 3 months differed from the healthy contralateral limb across all conditions tested. Peak isometric torque was reduced by 49% (P < 0.001). Peak isokinetic power was reduced by 29% at 30 degrees per second (P = 0.001), 24% at 120 degrees per second (P = 0.01), and 25% at 210 degrees per second (P = 0.01).

**Figure 5.**
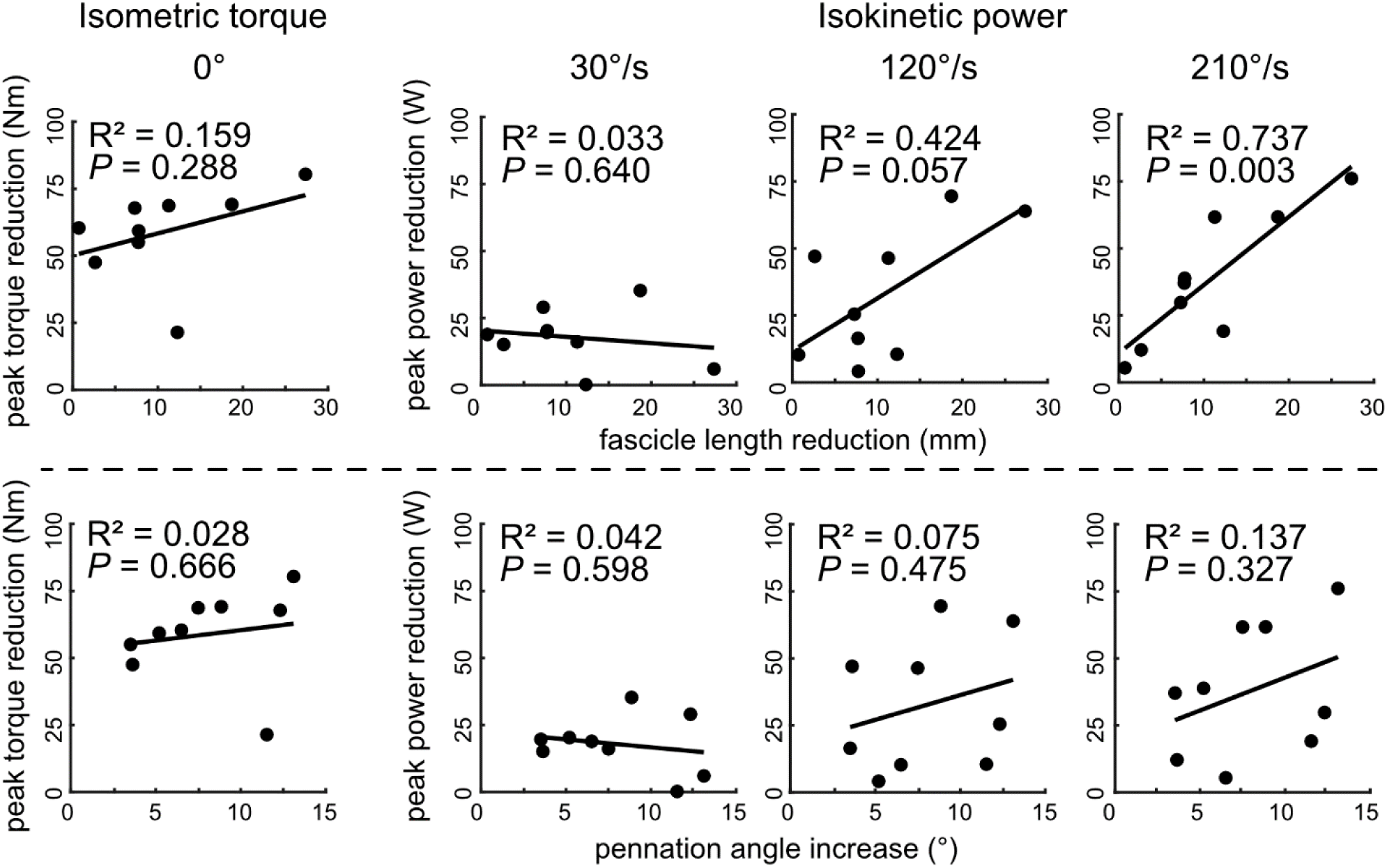
Deficits in peak torque during isometric plantarflexor contractions (*left column*) at neutral ankle angle and peak power during isokinetic plantarflexion contractions (*three columns on right*) throughout each patient’s ankle range of motion were correlated with decreases in fascicle length (*top*) and increases in pennation angle (*bottom*). Peak power deficits were moderately correlated with reductions in fascicle length at 120 degrees per second and strongly correlated with reductions in fascicle length at 210 degrees per second.

## DISCUSSION

This study challenges the paradigm that Achilles tendon ruptures are isolated tendon injuries. To do this, we longitudinally measured muscle-tendon structure at each clinical visit from initial presentation to the three-month follow-up appointment in a cohort of patients treated nonsurgically. We found that medial gastrocnemius fascicles retract to a shorter and more pennate configuration and the Achilles tendon elongates following the initial injury. Three months following injury, muscle fascicles were 13-19% shorter and 16-19% more pennate compared to the contralateral muscle (**Figure 2 and 4**). The magnitude of fascicle length remodeling at three months was strongly correlated with the magnitude of power deficits during fast plantarflexor contractions. These findings, for the first time, show that changes in muscle configuration following Achilles tendon ruptures explains the functional deficits experienced by patients when cleared to return to activity.

Our measurements of muscle-tendon structure and isokinetic plantarflexor function compare favorably with previous reports. To protect the healing tendon during the first 10 weeks following injury, we imaged the muscle and tendon with the foot supported in plantarflexion. These measurements agreed favorably with previous reports of the gastrocnemius muscle in peak plantarflexion.^20^ As expected, these measurements produced shorter fascicles compared to previous measurements of the medial gastrocnemius under load in neutral ankle position.^21,22^ However, when we measured muscle structure 3 months following injury, we did not have load restrictions and fascicle length and pennation measurements of the uninjured limb is very similar to previous reports throughout ankle range of motion.^20,21^

Early changes to muscle configuration following rupture was consistent amongst patients during the first 2 weeks before becoming more variable throughout healing. These increases in fascicle length variability throughout healing matches increased tendon loading that is prescribed using orthopaedic boots that progress to more and more weight bearing throughout the first 10 weeks. During the first two weeks following rupture, all patients were placed in a plantarflexed cast and restricted from loading the injured foot. However, as patients transition to a walking boot, the loading biomechanics of the healing tendon varies based on a myriad of patient-specific factors including number of steps taken, body weight, fit of the walking boot, ankle constraint, and muscle activity.^23–25^

Plantarflexor functional deficits measured at three months following Achilles tendon rupture were explained by gastrocnemius remodeling. Based on our previous computer simulations that show the isolated effects of shorter gastrocnemius fascicle length on plantarflexor function,^9,14^ these experimental findings confirm the link between changes in muscle structure and function. A previous report correlated fascicle length with heel raise height and found that patient reported outcomes were strongly correlated with the amount of fascicle remodeling in patients following surgical repair.^13^ These findings highlight the importance of skeletal muscle architecture and support the need for understanding the mechanisms that govern muscle remodeling following tendon rupture.

Muscle fascicle and tendon length appear to be coupled and detrimentally affected by Achilles tendon ruptures. When treated nonsurgically, we found that the ruptured tendon heals in an elongated position while the muscle fascicles remodel into a shorter and more pennate configuration. Similar observations were reported in patients treated surgically 3-12 months following treatment.^10^ Clinical treatment is targeted at minimizing tendon elongation following rupture, which can be targeted with early ankle motion^5^ and avoiding immobilization in dorsiflexion.^26^ Conversely, high velocity training stimulates longer muscle fascicles in a bipedal bird model.^27^ However, both tendon elongation and fascicle remodeling appear to occur during the weeks following the initial injury, limiting the potential efficacy of restoring muscle structure after the tendon is fully healed in an elongated position.^6^ Therefore, we suggest that future research focus on the interplay between tendon elongation and muscle remodeling throughout healing to identify treatment options that mitigate detrimental changes to structure and function.

Several limitations affected this study. These findings are a subset of data collected at each clinical visit throughout the first 3 months of treatment in an on-going 1-year prospective study. While, we plan to collect 1-year follow-up data on these patients, the purpose of this study was to determine the relationship between muscle remodeling and functional deficits when patients are cleared to return to activity. Similar decreases in gastrocnemius fascicle length up to 12 months following rupture have been reported,^10^ suggesting that the changes in muscle structure are permanent. We decided to image the medial gastrocnemius because it has longer, less pennate fascicles that are better suited for generating power in ankle plantarflexion compared to the soleus.^28^ Patients were instructed to perform voluntary maximal effort plantarflexion contractions, but we did not elicit maximal contractions with electrical stimulation. Plantarflexor power was not correlated with fascicle length of the uninjured limb, which we have previously reported in healthy young adults.^8^ which may be explained by patient guarding. However, the side-to-side differences in plantarflexion power were strongly correlated with differences in fascicle length, suggesting that intra-patient contractility was similar between the injured and contralateral limbs.

In conclusion, Achilles tendon ruptures lead to shorter and more pennate muscle fascicles that explain functional deficits in patients 3-months following injury. Treating these injuries nonsurgically mitigates the risks of surgery but does not re-oppose the ends of the ruptured tendon, resulting in tendon elongation and shorter muscle fascicles at 3 months. Future work should establish the link between tendon loading and muscle-tendon structure throughout rehabilitation, which may lead to improved plantarflexor structure and function.

## Authors’ Contributions

KO and JB developed the study; TH collected the data; TH and JB analyzed and interpreted the data; TH and JB drafted the manuscript; TH, KO, and JB revised the intellectual content of the manuscript; TH, KO, and JB approved the final version of the manuscript; and TH, KO, JB agreed to be accountable for all aspects of the study.

## ACKNOWLEDGEMENTS

This work was supported by The Thomas B. McCabe and Jeannette E. Laws McCabe Fund. We thank Rena Mathew and Shilpa Donde for assistance with collecting data.

***Acknowledgements*** the Authors have no acknowledgements

